# Exosome-Mediated Bidirectional Signaling Regulates Glioma Stem-Cell Homeostasis and Radiation-Induced Plasticity

**DOI:** 10.64898/2026.09.02.748690

**Authors:** Linda Azizi, Ling He, Aparna Bhaduri, Linda M. Liau, Harley I. Kornblum, Frank Pajonk

## Abstract

**Background:** Glioblastoma (GBM) is an aggressive brain malignancy characterized by therapeutic resistance and frequent recurrence. Glioma-initiating cells (GICs), also known as glioma stem cells (GSCs), contribute to these features through self-renewal and resistance to genotoxic stress. Radiation therapy can paradoxically replenish the GIC compartment by inducing stem-like properties in non-stem glioma cells. We investigated whether exosomes mediate bidirectional communication that maintains the balance between GIC and non-stem populations at steady state and following irradiation.

**Methods:** Exosomes were isolated from GIC-enriched gliomaspheres and differentiated monolayer cultures of the patient-derived GBM lines HK-374 and HK-390. Vesicles were characterized by transmission electron microscopy, nanoparticle analysis, and detection of CD63. Recipient cells were exposed to exosomes with or without 4 Gy irradiation. GIC abundance and function were evaluated using a ZsGreen-ornithine decarboxylase degron reporter, sphere-formation assays, and extreme limiting dilution analysis. Exosomal protein cargo was characterized by liquid chromatography-tandem mass spectrometry and Gene Ontology enrichment analysis. Single-cell RNA sequencing was used to evaluate changes in cellular composition upon exosome treatment.

**Results:** Isolated vesicles displayed characteristic exosomal morphology, a mean diameter of 41.6 +/- 14.7 nm, and CD63 expression. Irradiation produced a 12-fold increase in GIC reporter-positive cells among initially reporter-negative differentiated cells. Gliomasphere-derived exosomes significantly and dose-dependently suppressed this radiation-induced phenotype conversion and reduced functional GIC frequency and self-renewal in both patient-derived lines. Conversely, exosomes from GIC-depleted monolayer cultures increased reporter-positive cells two- to three-fold and enhanced the frequency and self-renewal of existing GICs. Proteomic analysis identified 1,796 exosomal proteins, including 336 differentially abundant candidates. Gliomasphere-derived exosomes were enriched in proteins associated with translation, RNA binding, actin organization, cytoskeletal regulation, and intracellular trafficking. Monolayer-derived exosomes were enriched in extracellular-matrix and stem-cell-niche components, including NID2, LAMA5, LAMB1, LAMB2, LAMC1, THBS1, TGFBI, IQGAP3, and APOE. Single-cell transcriptomic analysis showed that gliomasphere-derived exosomes prevented the radiation-induced expansion of neural progenitor-like cells, consistent with suppression of an induced stem-like state. Treatment of gliomaspheres with monolayer-derived exosomes left their overall cellular composition largely but enriched MYC-target, E2F-target, and oxidative-phosphorylation programs, suggesting that increased sphere formation may partly reflect enhanced proliferative capacity.

**Conclusions:** GBM cells use exosomes to establish a bidirectional feedback circuit between stem-like and differentiated tumor-cell compartments. GIC-derived exosomes constrain radiation-induced conversion of non-stem glioma cells into GICs, whereas differentiated-cell-derived exosomes promote a niche that supports GIC maintenance and self-renewal, potentially through expansion of a mixed-vascular-like population containing neurovascular progenitors. Whether exosomal IQGAP3 contributes to the expansion of this GIC population remains to be determined. These findings identify exosome-mediated intercellular communication as a potential mechanism regulating GBM cellular homeostasis, treatment resistance, and recurrence.

**Graphical Abstract:** 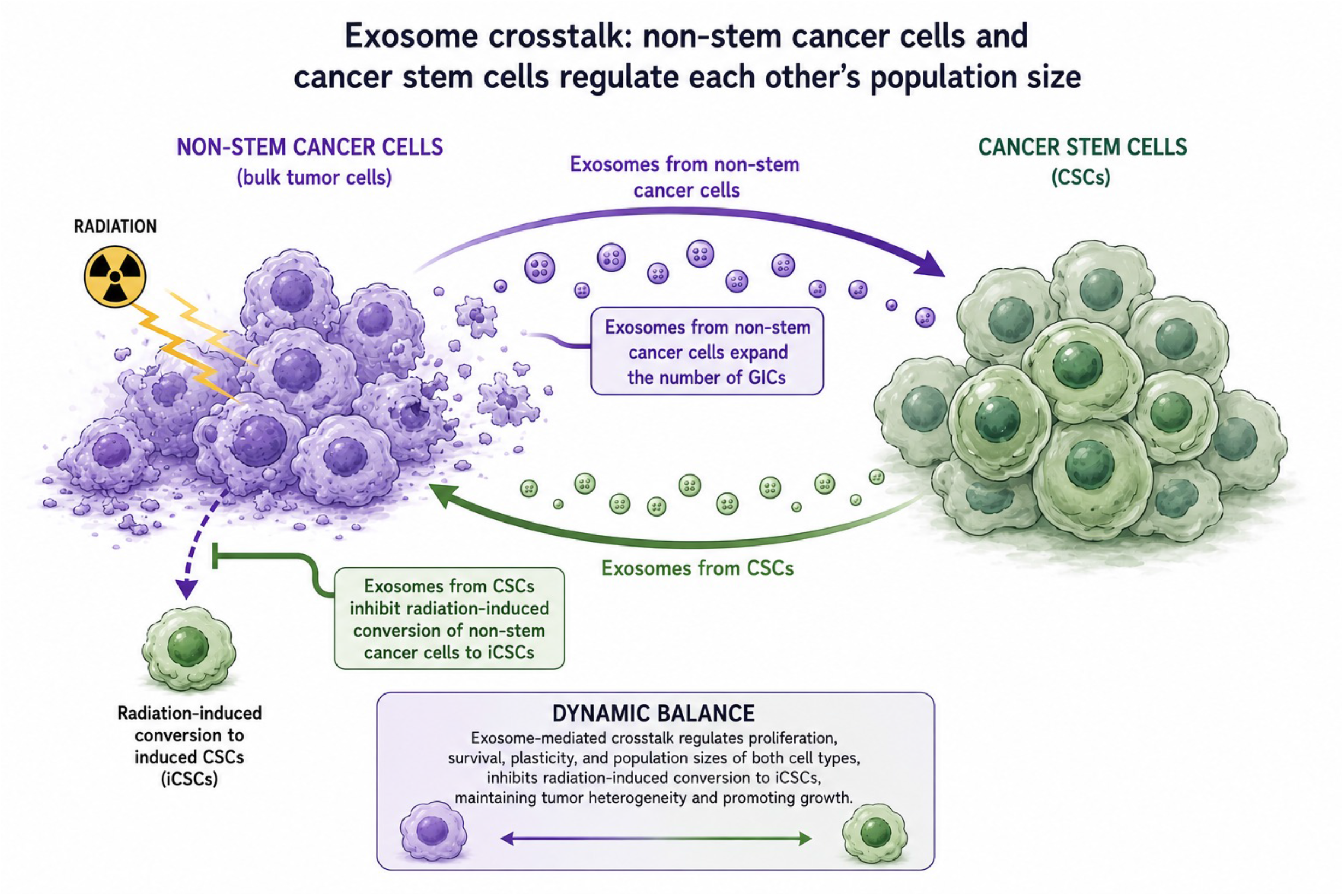

## Introduction

Glioblastoma (GBM) is the most aggressive primary brain malignancy in adults and is associated with a dismal prognosis. The current standard of care -surgical resection followed by radiation therapy (RT) with concurrent and adjuvant temozolomide-provides limited durable control, and tumor recurrence remains nearly inevitable [1], Treatment resistance and recurrence are attributed in part to a small population of highly resistant glioma-initiating cells (GICs), also known as glioma stem cells (GSCs) [2, 3]. Moreover, previous reports have shown that RT and chemotherapy can induce non-stem glioma cells to acquire GIC-like properties [3–5]. Because cellular plasticity is now recognized as a hallmark of cancer [4–7], defining the mechanisms that regulate transitions between GSC and non-stem states may reveal new therapeutic opportunities for limiting GBM heterogeneity, resistance, and recurrence.

Exosomes are nanosized membrane-bound extracellular vesicles, typically 20–150 nm in diameter, that mediate intercellular communication [8, 9]. Their cargo -including proteins, microRNAs, long noncoding RNAs, lipids, and metabolites-can influence tumor-cell growth, survival, and phenotype [8]. Signals from the tumor microenvironment can drive differentiated cancer cells to acquire stem-like properties, creating a dynamic and reversible equilibrium between stem-like and non-stem states [10]. Consistent with such feedback regulation, a previous study demonstrated that breast cancer stem cells suppress the RT-induced conversion of non-stem tumor cells into cancer stem cells [11].

Exosomes may help maintain this equilibrium by transmitting signals bidirectionally between stem-like and differentiated tumor-cell populations [10]. Here, we show that exosome-mediated signaling between GSCs and non-stem glioma cells regulates their relative abundance of these cellular compartments in GBM. This reciprocal communication establishes a bidirectional feedback mechanism that contributes to tumor-cell homeostasis at steady state and dynamically reshapes the balance between stem-like and non-stem populations following radiation-induced genotoxic stress.

## Materials and Methods

### Cell lines

Patient-derived xenograft (PDX) lines were established at UCLA as described in [12]. Characteristics of specific glioma sphere lines can be found in [13]. HK-374 and HK-390 cells (primary, IDH-wildtype, both classical TCGA-subtype) were cultured as gliomaspheres under serum-free conditions in ultra-low adhesion plates in DMEM/F12, supplemented with SM1 Neuronal Supplement (#05177, STEMCELL Technology, Kent, WA), EGF (#78006, STEMCELL Technology), bFGF (#78003, STEMCELL Technology), and heparin (1,000 USP Units/mL, NDC0409-2720-31, Lake Forest, IL) as described previously [12–14]. For marker-based GSC studies, cells were transduced with a lentiviral vector expressing the fluorescent protein ZsGreen, fused to the C-terminal degron of murine ornithine decarboxylase (ZsGreen-cODC), which reports the absence of proteasome activity in putative GSCs through accumulation of ZsGreen-cODC [15]. Monolayer cultures of both lines were grown in log-phase in DMEM supplemented with 10% exosome-depleted fetal bovine serum (Thermo Fisher Scientific, MA), 1 % penicillin, and streptomycin. All cells were grown in a humidified atmosphere at 37°C with 5% CO_2_. The identity of all patient-derived specimens was confirmed by DNA fingerprinting (Laragen, Culver City, CA). Cell lines were routinely tested for mycoplasma infection (#G238, Applied Biological Materials, Ferndale, WA).

### Exosome Isolation

Cell culture media from both patient-derived HK-374 and HK-390 glioma spheres and monolayers were collected when they reached 80% confluency. Extracellular vesicles (EVs) were obtained using the Total Exosome Isolation ™ cell culture media reagent (Invitrogen/Thermo Fisher Scientific, MA) following the manufacturer’s protocol or via ultracentrifugation. A starting volume of 240 ml of each cell culture medium was used to isolate the exosomes. For ultracentrifugation, exosome-depleted cell culture media were collected after 72 h of incubation and centrifuged at 300 x *g* for 10 min. The supernatant was collected and centrifuged at 2,000 x *g* for 10 min, followed by Ultracentrifugation (Beckman Coulter Life Sciences, USA) at 10,000 x *g* for 30 min. Supernatant was further centrifuged at 120,000 x *g* for 90 min. The pellet was washed in a large volume of phosphate-buffered saline (PBS) to eliminate contamination of proteins and centrifuged again at 120,000 x *g* for 90 min. Pelleted exosomes were resuspended in PBS, aliquoted in Eppendorf tubes, and stored at -80 °C for further downstream analysis.

### Exosome and Protein Quantification

Exosome samples were quantified by total protein concentration using the Micro BCA Protein Assay Kit (Thermo Fisher Scientific, MA), following Minimal Information for Studies of Extracellular Vesicles (MISEV) guidelines and literature for assessing approximate EV abundance through protein quantification [16, 17]. The Micro BCA assay was chosen for its higher sensitivity in detecting low protein concentrations (0.5-20 µg/mL range), which is well-suited for exosome preparations that typically have lower yields than conventional protein samples. Isolated exosomes were lysed in ice-cold RIPA lysis buffer containing 10 mM Tris-HCl (pH 8.0), 1 mM EDTA, 1% Triton X-100, 0.1% sodium deoxycholate, 0.1% SDS, 140 mM NaCl, and 1 mM PMSF, supplemented with proteinase inhibitors (Thermo Fisher Scientific, MA). Protein quantification was performed using the Micro BCA Protein Assay Kit following the manufacturer’s instructions. Absorbance was read at 562 nm using a SpectraMax iD3 microplate reader (Molecular Devices, San Jose, CA, USA).

### Characterization of Exosomes

Isolated exosomes from HK-374 glioma spheres were resuspended in PBS. Before starting the negative staining procedure for Transmission Electron Microscopy (TEM), Formvar carbon 300 mesh copper grids (Ted Pella Inc., CA) were glow-discharged for 30 seconds. 2.5 µL of the exosome sample was added to the grid and after 1 minute, the excess sample was blotted using a filter paper Whatman, Grade 1 (Whatman PLC, UK).12.5 µL 2% uranyl acetate (UA) (Electron Microscopy Sciences, PA) was added immediately drop by drop, while blotting in between each drop and the last drop was left on the grid for 1 minute. After blotting the excess UA, the grid was dried at room temperature. Imaged the grid on a T12 TEM (FEI Tecnai, Hillsboro) operating at 120 kV. Vesicle diameters (nm) were measured using ImageJ and then imported into Microsoft Excel for downstream analysis. Particle sizes were grouped into predefined bins, and the frequency of particles per size bin was calculated. Exosomes were counted using a ViewSizer 3000 nanotracking particle size analyzer (HORIBA, Irvine, CA). 300ng/ml of exosomal protein corresponded to 3.89 ± 0.87 x 10^9^ particles/ml.

Whole cell lysates for both cell lines were prepared for Western blotting using ice-cold 1x Ripa buffer (10mM Tris-HCl (pH 8.0), 1 mM EDTA, 1% Triton X-100, 0.1% Sodium Deoxycholate, 0.1% SDS, 140 mM NaCl, 1 mM PMSF) containing proteinase inhibitor (Thermo Fisher Scientific, MA). Protein concentrations were determined using the BCA protein assay kit (Thermo Fisher Scientific). EV samples were diluted 1:1 with ice-cold 1X RIPA buffer, and protein concentrations were determined using the micro-BCA protein assay kit (Thermo Fisher Scientific, MA). For western blot analysis of tetraspanin marker CD63, samples were prepared under non-reducing conditions. This approach was necessary because the epitopes recognized by antibodies against these proteins are dependent on intact disulfide bonds [18]. Therefore, no reducing agent was added to the 4x Laemmli sample buffer (Bio-Rad Laboratories Inc., CA) before electrophoresis. Samples were then heated at 95°C for 10 minutes, loaded onto 10% SDS-PAGE gels, and subjected to electrophoresis for 2h. Samples were then transferred onto a nitrocellulose membrane (Bio-Rad Laboratories Inc., CA) and blocked with 5% bovine serum albumin (BSA) in 1x TBST for 30 minutes at room temperature, followed by incubation with primary antibodies against CD63 (Rabbit mAB #52090, 1:1000, Cell Signaling Technology, MA) in 5% BSA overnight at 4°C with gentle rocking. Membranes were then washed three times for 5 minutes each with 1X TBST and incubated with a secondary antibody (anti-rabbit, horseradish peroxidase-linked #7074,1:1000, Cell Signaling Technology) in 1X TBST for two hours at room temperature and gentle rocking. Membranes were washed again three times for 5 minutes each with 1X TBST. For chemiluminescent western blot analysis, Prometheus Pico ECL spray (Genesee Scientific, NC) was added to each membrane and incubated at room temperature for 5 minutes. The blots were then scanned using the Odyssey FC imager (LI-COR Biosciences, Lincoln, NE).

### Irradiation

Cells were irradiated at a single dose of 4 Gy at room temperature using an X-ray irradiator (Gulmay Medical Inc., Atlanta, GA) at a dose rate of 5.519 Gy/min. Control samples were sham-irradiated. The X-ray beam was operated at 300 kV and hardened using a 4 mm Be, a 3 mm Al, and a 1.5 mm Cu filter and calibrated using NIST-traceable dosimetry.

### Sphere Formation Assay (SFA) and Extreme Limiting Dilution Analysis (ELDA)

To assess the effect of exosomes on self-renewal monolayer cultures were depleted of putative GSCs (ZsGreen-cODC-positive) by FACS as described previously [15]. Monolayer cultures treated with exosomes isolated from glioma spheres were trypsinized/digested on day 5 after exosome treatment, and cells were serially diluted in non-tissue culture-treated 96-well plates under serum-free conditions at a range from 1 to 512 cells/well. Growth factors (EGF and bFGF) were supplemented every two days. Glioma spheres were counted 7 days later and presented as a percentage of the initial number of cells plated. GSC frequencies were calculated using the ELDA software package in R [19].To further investigate the bidirectional communication between glioma stem cells and non-stem glioma cells, we designed a “reverse experiment” by culturing HK-374 and HK-390 glioma spheres, which are enriched for (GSCs), in non-adherent 6-well plates, and this time treated the glioma spheres with isolated exosomes from monolayers, which are mostly differentiated. Based on dose response ELDA experiments, we chose an exosome concentration of 300 ng/ml for consecutive experiments.

### Migration Assay

HK-390 or HK-374 monolayers were plated on 3 cm Petri dishes in 10% exosomes-depleted FBS medium and treated with or without 300 ng/ml exosomes isolated from HK-390 or HK-374 glioma spheres in combination with a single dose of RT at 4 Gy. 48 hours post-RT, the monolayers were serum starved in 1% exosome-depleted FBS medium. The next day, cells were trypsinized and plated on cell culture inserts with an 8 µm pore size (Corning Inc., NY) at 100K cells/well. To facilitate cell migration in the chambers, an FBS gradient was created by filling the bottom chambers with 10% exosomes-depleted FBS medium. After 20-24 hours of incubation, cells were fixed using 10% formalin and washed with PBS. Using a cotton swab, non-migrated cells on the upper side of the membrane were gently removed, and migrated/attached cells at the bottom of the membrane were stained with 1% crystal violet. Subsequently, images were taken with a digital microscope (BZ-9000, Keyence, Itasca, IL), and migrated cells were counted using the Image J software.

### Mass Spectrometry

#### In-solution digestion

Exosome samples from both HK-374 glioma spheres and monolayers were centrifuged, and the pellets were resuspended with 50 µl 8M Urea followed by sonication. Samples were then incubated for 15 min at -80 °C for further lysis. Proteins in solution (50 μL) were reduced by 5 mM TCEP at 56 °C for 1 hr and alkylated by 40 mM iodoacetamide at room temperature for 30 min in the dark. Trypsin dissolved in 50 mM ammonium bicarbonate was added to protein pellets for digestion at 37°C overnight. Protein digests were desalted using Empore stage-tips the following day. The elution from the stage-tip was dried by speed vac and re-suspended in 3% acetonitrile with 0.1% formic acid.

#### LC MS/MS

1.0 µg protein was injected into an UltiMate 3000 nanoLC, equipped with a 75 µm x 2 cm trap column packed with C18 3 µm bulk resins (Acclaim PepMap 100, Thermo Scientific, MA) and a 75 µm x 15 cm analytical column with C18 2 µm resins (Acclaim PepMap RSLC, Thermo Scientific). The nanoLC gradient was 3−35% solvent B (A = H2O with 0.1% formic acid; B = acetonitrile with 0.1% formic acid) over 40 min and from 35% to 85% solvent B in 5 min at a flow rate of 300 nL/min. The nanoLC was coupled to a Q Exactive Orbitrap mass spectrometer (Thermo Fisher Scientific, MA). The ESI voltage was set to 1.9 kV, and the capillary temperature was set to 275° C. Full spectra (m/z 350 - 2000) were acquired in profile mode with a resolution of 70,000 at m/z 200 and an automated gain control target of 3 × 10^6^. The most abundant 15 ions were subjected to fragmentation by higher-energy collisional dissociation (HCD) with normalized collisional energy of 25. MS/MS spectra were acquired in centroid mode with a resolution of 17,500 at m/z 200. The AGC target for fragment ions was set to 2 × 10^4^ with a maximum injection time of 50 ms. Charge states 1, 7, 8, and ‘unassigned’ were excluded from tandem MS experiments. Dynamic exclusion was set to 45.0 s.

#### Data Analysis

Raw data was searched against the Uniprot human database by Proteome Discoverer (version 2.5) for protein identification. The following parameters were set: precursor mass tolerance ±10 ppm, fragment mass tolerance ±0.02 Th for HCD, up to two mis-cleavages by semi trypsin, methionine oxidation as a variable modification, and cysteine carbamidomethylation as a static modification.

### Protein Bioinformatics Analysis

Mass spectrometry analysis identified 1796 proteins in exosomes isolated from both and monolayer samples. Proteomic data were filtered to include proteins with an FDR < 1% and a significance threshold of p < 0.05. Proteins were then ranked by their abundance ratio (NS-Exo’s/Mono-Exo’s), and those with ratios ≥ 2.0 were selected as significantly enriched in the neurosphere population. Conversely, proteins with NS/Mono ratios < 1.0, representing those upregulated in Mono-Exo’s, were chosen for comparative analysis. The resulting protein datasets from both Mono- and NS-Exo’s samples were then used to generate a clustered hierarchical heatmap using RStudio (version 2025.05.1+513).

Top candidates were compared to the ExoCarta database of validated exosomal proteins, and Venn diagrams were generated using RStudio (version 2025.05.1+513) to visualize the overlap.

Gene Ontology (GO) enrichment analysis and gene-term network plots were generated using the R packages clusterProfiler, org.Hs.eg.db, AnnotationDbi, and enrichplot with default parameters. The top 20 most significantly enriched terms from each GO category (Ranked by FDR) were selected for visualization using RStudio (version 2025.05.1+513).

### Gene level Heat Maps

Protein abundance values were log2-transformed after addition of a pseudocount of 1. To account for differences in absolute abundance between proteins, values were standardized independently for each protein across the six samples (three monolayer and three gliomasphere samples) using row-wise Z-scores. The standardized abundance values were visualized as heat maps, with proteins arranged according to their assigned primary functional categories. Thus, heat-map values represent the relative abundance of each protein across experimental conditions rather than differences in absolute abundance between proteins. Positive Z-scores indicate abundance above the protein-specific mean, whereas negative Z-scores indicate abundance below the protein-specific mean.

### Functional Category Heat Map

Protein abundance values were log2-transformed after addition of a pseudocount of 1. To enable comparison of relative abundance patterns across proteins with different absolute abundance levels, values were standardized independently for each protein across the six samples (three monolayer and three gliomasphere samples) using row-wise Z-scores. Proteins were assigned to functional categories based on their primary biological function. For each category, the mean Z-score of all constituent proteins was calculated separately for each sample and visualized as a heat map. Positive and negative values therefore represent relative enrichment and depletion, respectively, compared with the mean abundance of each protein across all samples.

### Single Cell RNA Sequencing

Gliomaspheres in the suspension culture were collected and dissociated with TrypLE (no phenol red, Thermo Fisher Scientific), and the monolayer cells were washed with HBSS and de-attached with Trypsin/EDTA (Cell Applications, Cat#090K). The cells were then pooled and filtered through a 40-µm strainer and fixed with Evercode^TM^ Cell Fixation v3 kit (#ECF2101, Parse Biosciences, Seattle, WA) following the manufacturer’s guidelines. The samples were then sent out to the Genomics High Throughput Facility (GHTF) at the University of California, Irvine for subsequent single cell RNA sequencing (scRNAseq) using the Evercode^TM^ Whole Transcriptome Mini kit v3 (#EC-W01010, Parse Biosciences). Sequencing reads from the mRNA libraries were mapped to the human genome (hg38) using the Parse Trailmaker pipeline and filtered matrices were downloaded and processed using the Seurat R package. The following steps were performed: 1. Filtering out low-quality cells and genes. 2. Removal of doublets. 3. Normalization and scaling. 4. Dimensionality reduction using PCA. 5. and UMAP. 6. Cell type annotation in Seurat was performed using a recently published GBM meta-atlas [20]. Gene set enrichment analysis (GSEA) was performed using genes ranked by the differential-expression statistic obtained from the edgeR pseudobulk analysis. Pathways were defined using the Molecular Signatures Database (MSigDB) Hallmark gene-set collection. Enrichment scores were calculated across the complete ranked gene list, and multiple-testing correction was applied using the Benjamini–Hochberg method. Hallmark pathways with a false discovery rate (FDR) below [0.05] were considered significantly enriched.

### Statistics

Unless stated otherwise all data shown are represented as mean ± standard error mean (SEM) of at least 3 biologically independent experiments. A *p*-value of ≤0.05 in an unpaired two-sided *t*-test or one-way ANOVA for multiple testing indicated a statistically significant difference.

## Results

### Characterization of isolated EVs

EVs are heterogeneous in size and consist of microvesicles (100-1000 nm) and exosomes (20-150 nm) [8, 9, 21]. Characterizing EVs in this study using transmission electron microscopy revealed vesicles with the exosomal morphology: a typical lipid bilayer (**Figure 1A**) and diameters in the expected small EV range of 20-80 nm (**Figure 1B**) with a mean particle size of 41.6 ± 14.7 nm. Larger vesicles (>100 nm) were rarely observed, consistent with an enrichment for exosomes within our EV yield. Western blots confirmed the presence of the exosomal marker CD63 [22, 23] in the cell lysates of HK-374 and HK-390 and exosomes derived from both lines, appearing as a typical wide smear resulting from heavy glycosylation of CD63 (**Figure 1C**) [24, 25].

**Figure 1.**
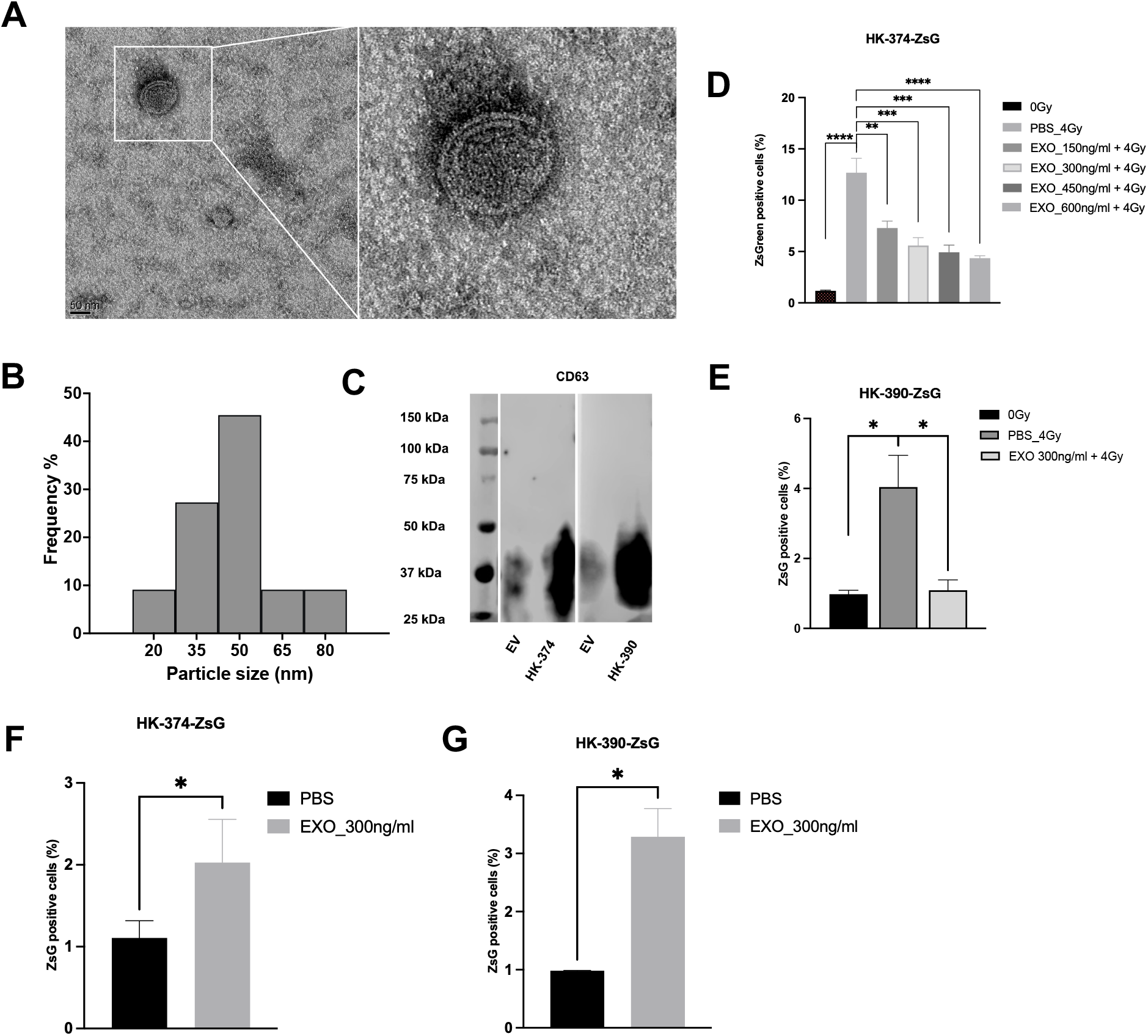
Validation of isolated exosomes. **A**) TEM picture of isolated exosomes shows the typical bi-lipid membrane. **B**) Size distribution of isolated exosomes analyzed with TEM and Image J, with a mean particle size of 41.63 ± 14.67 nm. **C**) Western blot of exosomal marker CD63 present in both HK-374 & HK-390 glioma spheres and their corresponding isolated exosomes. **D**) Treatment of HK-374-ZsGreen-cODC cells with 4 Gy leads to an increase in ZsGreen-cODC-positive cells and gliomasphere-derived exosomes inhibit this radiation-induced increase in a dose-dependent manner. **E**) Treatment of HK-390-ZsGreen-cODC cells with gliomasphere-derived exosomes at 300 ng/ml inhibit the radiation-induced increase in ZsGreen-cODC positive cells. All experiments represent at least 3 independent biological repeats. *P*-values were calculated using one-way ANOVA for D and E; unpaired student t-test for F and G. * *p*-value < 0.05, ** *p*-value < 0.01, *** *p*-value < 0.001, **** *p*-value < 0.0001.

### Exosomes derived from glioma stem cells inhibit the self-renewal of existing stem cells

To investigate if exosomes modulate the ratio of GICs to non-stem glioma cells at steady state or in response to radiation, we performed phenotype conversion assays using a fluorescent reporter system for GICs [26]. It is based on the stable expression of the fluorescent protein ZsGreen linked to the C-terminal degron of ornithine decarboxylase that channels ZsGreen to immediate, ubiquitin-independent proteasomal degradation [27]. In cells lacking proteasome activity, ZsGreen accumulates and these ZsGreen-positive cell populations are enriched for GICs [26]. Patient-derived HK-374-ZsGreen were sorted for ZsGreen-negative cells, plated as monolayers, irradiated with a single dose of 4 Gy on day 1 and exposed daily to different amounts of exosomes isolated from HK-374-ZsGreen GSC-enriched gliomaspheres. Controls were exposed to PBS and sham irradiated. On day five after irradiation cells were probed for ZsGreen-positive cells by flow cytometry. Irradiation with 4 Gy led to a significant, 12-fold increase in ZsGreen-positive cells compared to unirradiated control cells, which agreed with studies [26]. Treatment with gliomasphere-derived exosomes led to a significant, dose-dependent reduction in the percentage of radiation-induced ZsGreen-positive cells (**Figure 1D**). These findings were confirmed in a second patient-derived line, HK-390-ZsGreen (**Figure 1E**).

Next, we tested if exosomes derived from GIC-depleted monolayer cells would alter the proportion of GIC marker-positive cells in gliomaspheres. Using HK-374-ZsGreen and HK-390-ZsGreen cell lines we found that treatment of gliomaspheres with exosomes from GIC-depleted cells significantly increased the number of ZsGreen-positive cells 2- and 3-fold, respectively (**Figure 1F/G**). Together, these data suggested that non-stem glioma cell populations and glioma-initiating cell populations regulate each other’s population size via exosomes-mediated signaling.

GICs are not defined by the expression GIC markers but only by the presence of functional cancer stem cells traits, including self-renewal. To assess if exosomes treatment would affect the proportion of functional GICs in response to radiation, HK-374 cells were plated at as monolayers, irradiated, treated with exosomes for five consecutive days and then re-plated at clonal densities under sphere-forming conditions. Seven days later, sphere-formation was assessed and the data subjected to an extreme limiting dilution analysis (ELDA) [19]. The addition of exosomes significantly reduced glioma stem cell frequencies in a dose-dependent manner, thus compromising self-renewal capacity (**Figure 2A/B**). While all exosome doses showed significant biological activity, subsequent experiments were conducted using 300 ng/ml of exosomes, a dose at which the effect of exosomes plateaued.

**Figure 2.**
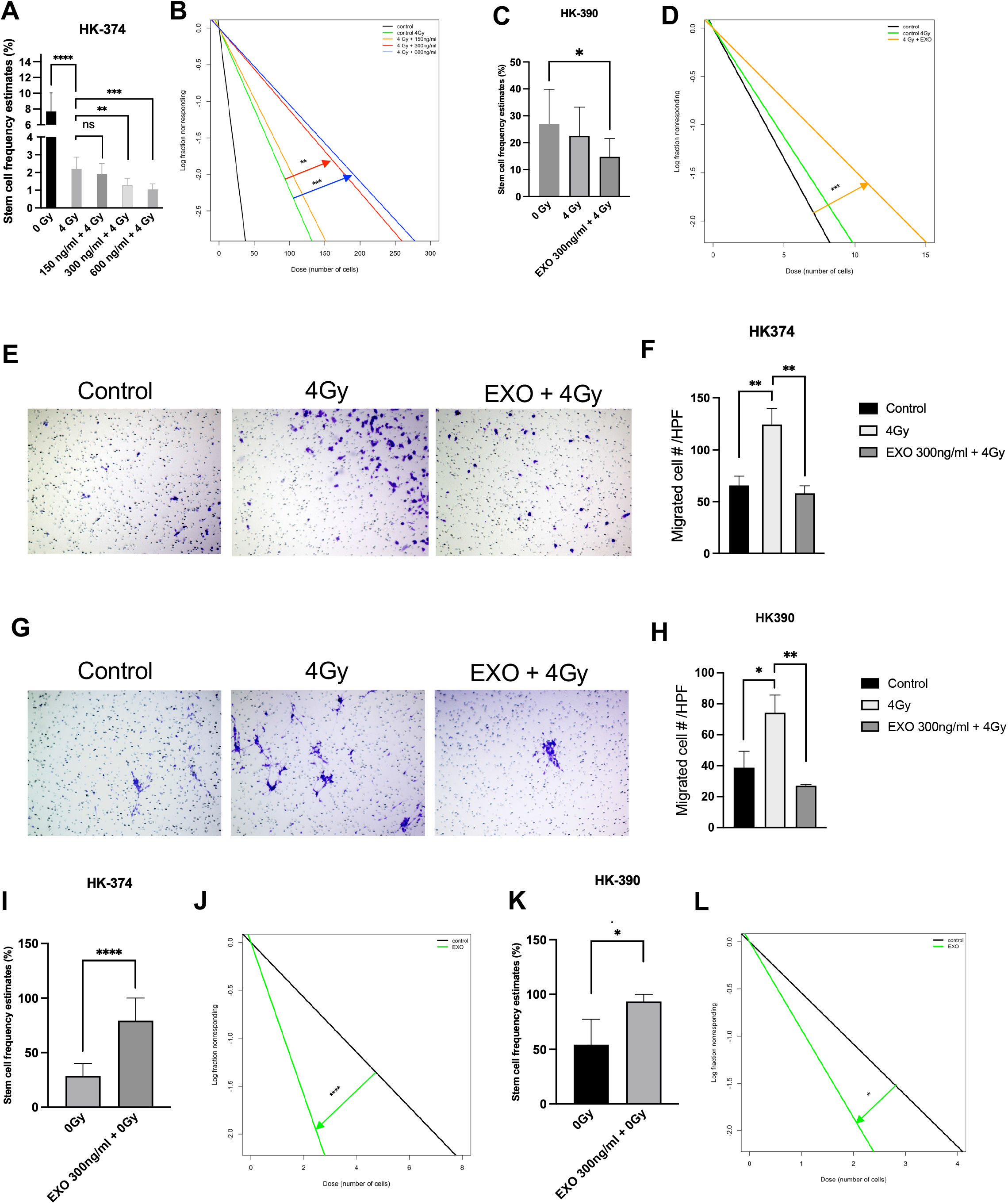
Effects of exosomes on GSC populations. **A/B**) HK-374 or **C/D**) HK-390 monolayers were irradiated with a single dose of 4Gy and treated with different concentrations of isolated exosomes from HK-374 or HK-390 gliomaspheres. Assessed by ELDAs, exosome treatment reduced GSC frequencies and impaired self-renewal. A single dose of 4 Gy significantly increased the migration of HK-374 **E/F**) or HK-390 cells **G/H**). This increase in migration was significantly reduced by treatment with exosomes from corresponding gliomaspheres. HK-374 **I/J**) or HK-390 **K**/L) gliomaspheres were exposed to 300 ng/ml of isolated exosomes from HK-374 or HK-390 monolayers, respectively, subjected to sphere-formation assays, and analyzed via ELDA. Exosome treatment significantly increased the GSC frequencies and promoted self-renewal. All experiments represent at least 3 independent biological repeats. Stem cell frequencies and calculated statistics were obtained using the ELDA software [19]. Data are shown as mean 95% confidence intervals. *P*-values were calculated using one-way ANOVA for F and H. * *p*-value < 0.05, ** *p*-value < 0.01, *** *p*-value < 0.001, **** *p*-value < 0.0001.

To validate our findings, we applied the same treatment protocol to an additional patient-derived GBM cell line, HK-390. Daily treatment of HK-390 cells with exosomes derived from corresponding gliomaspheres led to compromised self-renewal capacity with a comparable decrease in stem cell frequencies (**Figure 2C/D**).

Next, we tested if the same treatment with exosomes would affect radiation-induced migration of glioma cells. A single dose of 4 Gy led to a significant increase in the number of migrating HK-374 (**Figure 2E/F**) or HK-390 cells (**Figure 2G/H**) and this was significantly reduced by treatment with exosomes derived from corresponding glioma spheres.

### Exosomes derived from non-stem glioma cells promote self-renewal of existing glioma stem cells

Next, we sought to test if exosomes from monolayer cultures, depleted from GICs, would affect the size of the GIC cell population in gliomaspheres. HK-374 and HK-390 gliomaspheres were treated daily with exosomes at 300 ng/mL. On day 5 of treatment HK-374 (**Figure 2I/J**) and HK-390 (**Figure 2K/L)** glioma spheres were digested into single cell suspensions and reseeded at clonal densities. Seven days later, sphere numbers were assessed and subjected to ELDAs. Exosome treatment led to a significant increase in glioma-initiating cell frequencies compared to untreated controls, indicating a gain in self-renewal capacity.

### Cargo of GIC-vs. non-stem cell-derived exosomes show distinct abundance pattern

To characterize the differential protein enrichment between non-stem and stem cell populations in GBM, we employed mass spectrometry and proteomic analysis to identify the protein cargo of exosomes samples isolated from HK-374 monolayers, primarily representing non-stem glioma cells, and gliomaspheres, enriched for glioma-initiating cells. To validate the exosomal origin of the identified proteins in mass spectrometry, a total of 336 differentially abundant cargo proteins (DAPs) were compared to the ExoCarta database, a comprehensive resource of exosomal proteins [28–30]. 322 proteins out of the 336 top candidates were present in ExoCarta (**Supplemental Figure 1A**). Only 14 proteins present in exosomes from gliomaspheres had not previously been known as exosome cargo. A list of all identified proteins is given in **Suppl Table 1.** DAPs for gliomasphere- and monolayer-derived exosomes samples were hierarchically clustered and visualized in a heatmap (**Supplemental Figure 1B**), revealing distinct cargo patterns. The presence of the exosomal markers CD9, CD81, CD63, TSG101, HSP70 and the absence of the non-exosomal marker GOLGA2 [31] further indicated the purity of exosome preparations (**Supplemental Figure 1C**).

Next, we performed a Gene Ontology (GO) enrichment analysis for gliomasphere- and monolayer-derived DAPs. The top 20 enriched GO terms for the DAPs from gliomasphere- and monolayer-derived exosomes are shown in **Figure 3A/B**. The top 20 enriched GO terms for the 319 DAPs from gliomasphere-derived exosomes mainly covered Biological Processes of actin filament organization, polymerization, depolarization and caping, actin nucleation as well as cytoplasmic translation. Gene level and functional category heat maps are shown for actin filament organization (**Figure 3C/D**) and for cytoplasmic translation (**Figure 3E/F**). The top 20 enriched GO terms for the 17 DAPs from monolayer-derived exosomes mainly covered Biological Processes of extracellular matrix, structure, and basement membrane organization, cell adhesion and proliferation, and integrin signaling (**Figure 3G/H**). The full list of GO terms is given in **Supplementary Tables 2 and 3**.

**Figure 3.**
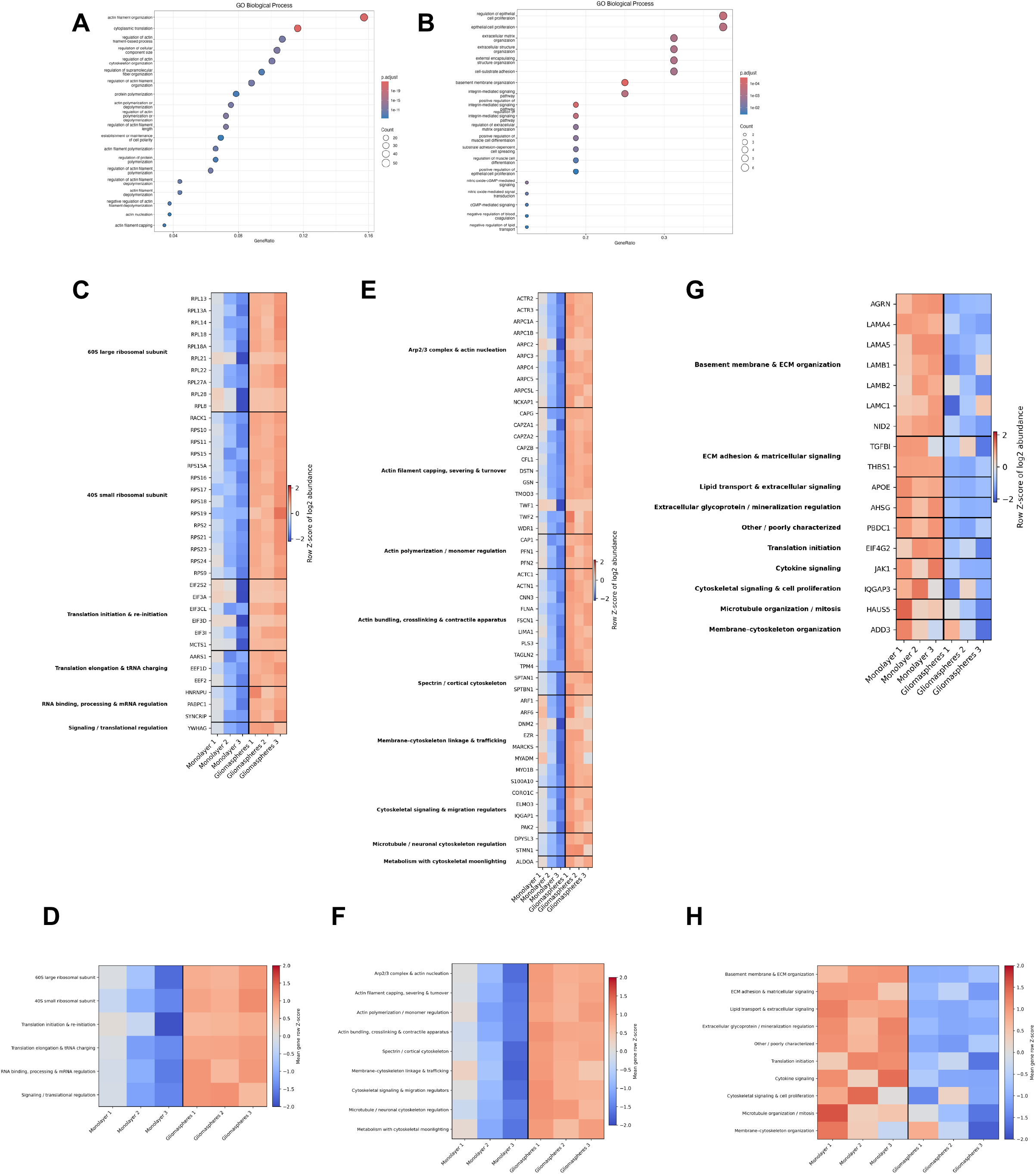
Mass spec analysis of exosome populations isolated from gliomaspheres and monolayers of HK-374 cells. **A**) Gene ontology (GO) terms of top 20 biological processes enriched in gliomasphere- and **B**) non-stem glioma cell-derived exosomes. Gene level heat maps for GO terms *Cytoplasmic translation* (**C/D**) and *Actin Filament Organization* (**E/F**) for DAPs in gliomasphere-derived exosomes and all 17 DAPs in monolayer-derived exosomes (**G/H**). For gene level heat maps (**C/E/G**) protein abundance values were log2-transformed and standardized independently for each protein across the six samples using row-wise Z-scores. The resulting Z-scores were visualized as a heat map, with proteins arranged according to their assigned functional categories. Positive and negative values indicate relative enrichment and depletion, respectively, compared with the mean abundance of each protein across all samples. For functional category heat maps (**D/F/H**) protein abundance values were log2-transformed and standardized for each protein across the six samples using row-wise Z-scores. Proteins were grouped by functional category, and the mean Z-score of all proteins within each category was calculated for each sample and visualized as a heat map. Positive and negative values indicate relative enrichment and depletion, respectively.

### Exosome cargo bi-directionally affects GSC populations at steady-state and after irradiation

Finally, we performed single cell RNA sequencing (**Figure 4A**) to test if exosomes caused changes in cell type composition that would link exosome cargo to changes in the proportions of functional GICs. Cell annotation using a previously published cell meta-atlas for GBM (**Figure 4B**) revealed a lack of complexity of differentiated HK-374 cells, with mesenchymal- and mixed vascular-like dominating the cell type composition (**Figure 4C/D**). A single dose of 4 Gy doubled the proportion of mesenchymal-like cells reduced the proportion of mixed-vascular-like and cycling cells, with the latter agreeing with the well-known radiation-induced cell cycle arrest. These population changes were not affected by treatment with exosomes obtained from GIC-enriched gliomasphere cultures. Radiation caused an increase in NPC-like cells, and this was prevented by exosome treatment (**Figure 4E/F**).

**Figure 4.**
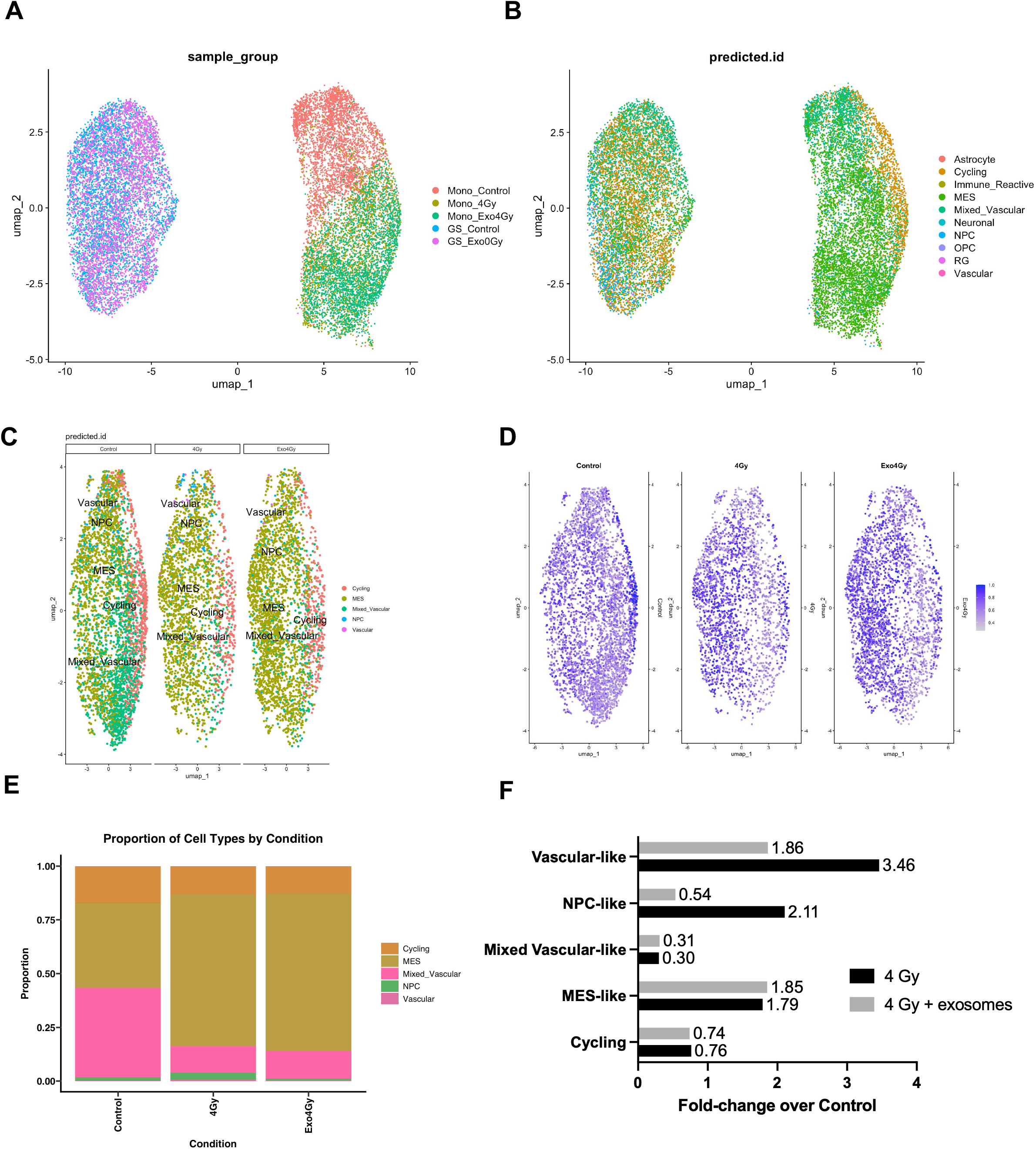
Single cell RNA sequencing of HK-374 monolayer cells. **A**) Dimensional reduction using UMAP for monolayer and gliomasphere cells. **B**) Annotation of cell types. **C**) Effects of radiation (4 Gy) and exosomes on cell types in monolayers. **D**) Predicted ID scores for cell type annotations. **E/F**) Radiation causes a 2-fold increase in NPC-like cells and this increase in prevented by treatment with gliomasphere-derived exosomes.

Gliomaspheres obtained from the same PDX line showed a higher level of heterogeneity with cycling cells, mesenchymal- and mixed-vascular-like cells as the most prominent populations (**Figure 5A/B**). Treatment with exosomes derived from differentiating cells decreased the proportion of all cell types in general, except for mixed vascular-like and astrocyte-like cells with the latter representing only 0.2 % of the total population. The proportion of mixed vascular-like cells only moderately increased 1.13-fold after exosome treatment (**Figure 5C/D**).

**Figure 5.**
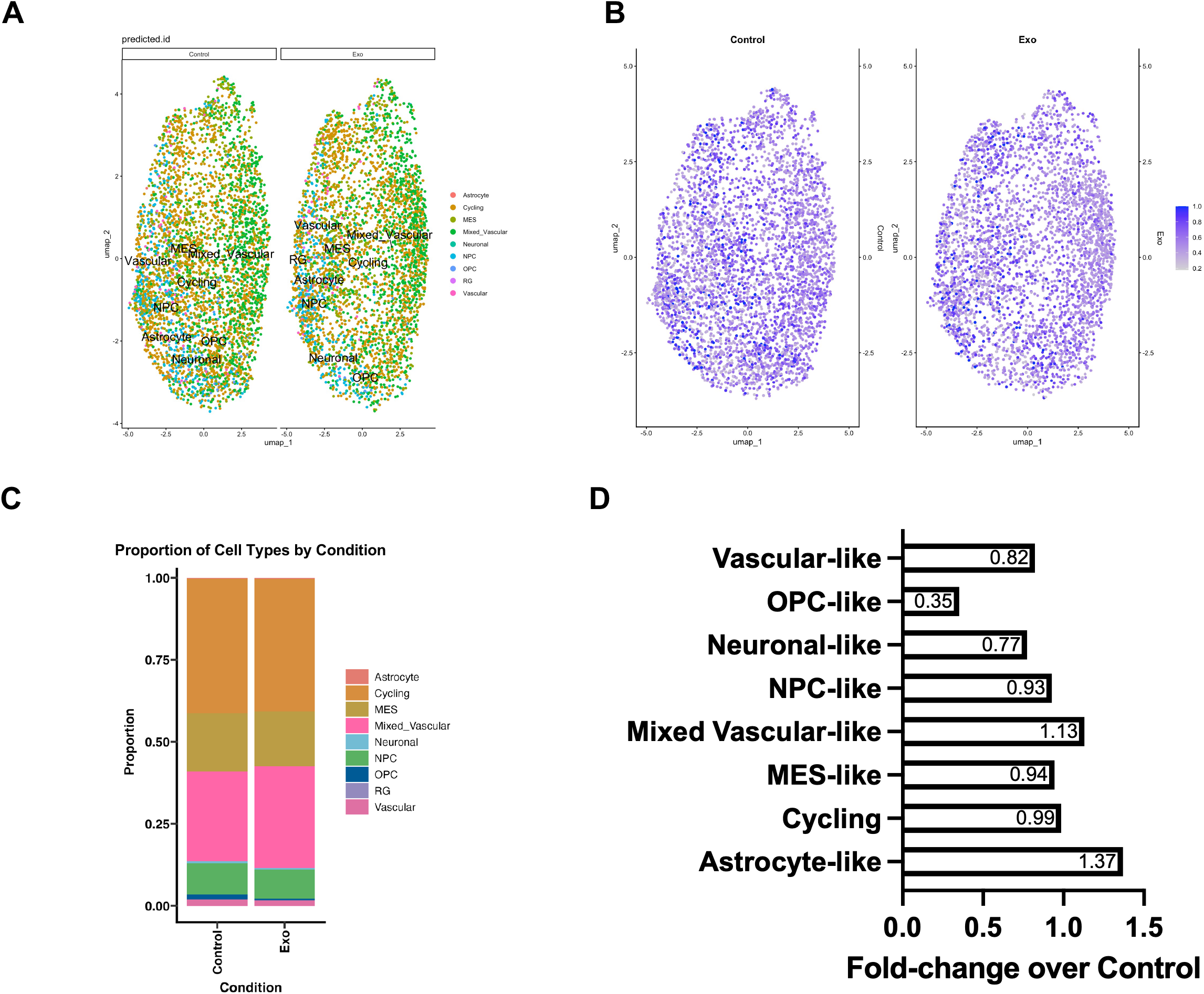
Single cell RNA sequencing of HK-374 gliomasphere cells. **A**) Effects of radiation (4 Gy) and exosomes on cell types in monolayers. **B**) Predicted ID scores for cell type annotations. **C/D**) Treatment of gliomaspheres with exomes from monolayers cells causes a 1.13-fold increase in mixed vascular-like cells.

GSEA of the scRNAseq data revealed positive enrichment of HALLMARK_MYC_TARGETS_V1 in the exosome-treated gliomaspheres for mixed vascular-like cells, cycling cells, mesenchymal-like cells, and NPC-like cells. Positive enrichment for HALLMARK_E2F_TARGETS was observed in mixed vascular-like cells, mesenchymal-like cells, and NPC-like cells. Positive enrichment for HALLMARK_OXIDATIVE_PHOSPHORYLATION was found in mixed vascular-like cells and cycling cells (**Figure 6**). Lists of the top 10 enriched gene sets are provided in the **Supplementary Table 3**

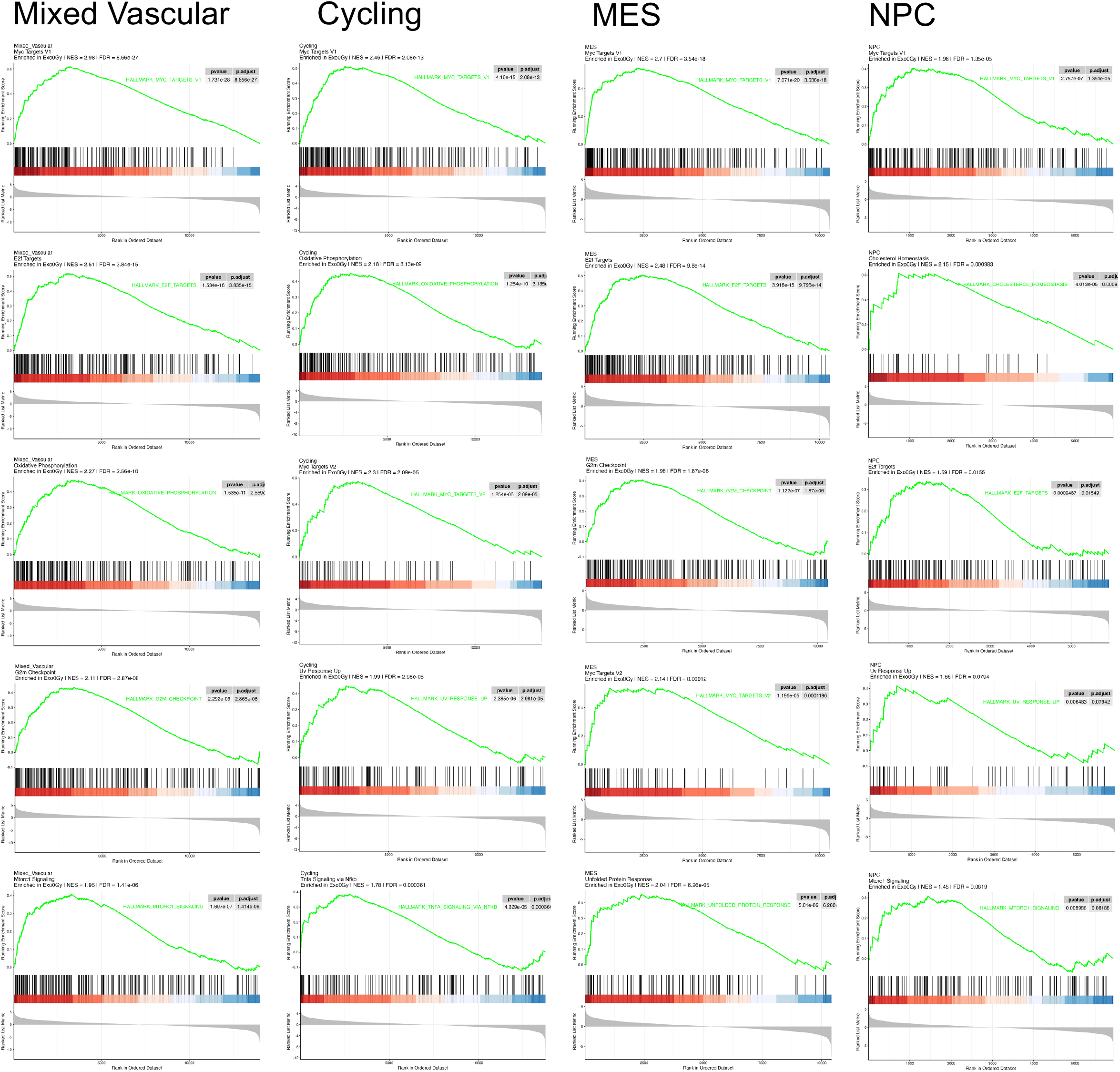

## Discussion

GBM is characterized by its highly aggressive nature and dismal prognosis, ranking among the deadliest brain cancers. The current standard of care, which involves surgical resection followed by RT with concurrent and adjuvant temozolomide, often results in high rates of tumor recurrence and treatment failure. Specifically, the presence of a small yet highly resistant population of glioma-initiating cells (GICs), also known as glioma stem cells (GSCs), is thought to underlie both treatment resistance and the high rate of tumor recurrence observed in patients with GBM [2, 3].

Paradoxically, RT and chemotherapy can further replenish the GIC pool by converting non-GICs into induced GICs through epigenetic reprogramming, a process termed *RT-induced phenotype conversion*, thus predominantly eliminating but also promoting treatment resistance and tumor recurrence [2].

In the absence of external perturbations, hierarchically organized tissues rely on feedback mechanisms to regulate the balance among symmetric self-renewing, symmetric differentiating, and asymmetric stem cell divisions. This regulation compensates for cell loss, maintains occupancy of a limited number of stem cell niches, and preserves tissue homeostasis. In response to genotoxic stress and depletion of the stem cell compartment, the balance may shift toward symmetric self-renewal to restore the stem cell pool, returning toward steady-state dynamics once the compartment has been replenished. The similar principles apply to tumors, although the ratios of symmetric to asymmetric divisions are differentially regulated to support linear or exponential tumor growth.

The net effect of radiation on the CSC compartment reflects the balance between two opposing processes: radiation-induced cell killing and the acquisition of stem-like properties by non-stem cancer cells [11, 15]. At clinically relevant radiation doses, cell killing predominates, resulting in a net reduction in CSC numbers. However, radiation-induced phenotypic conversion of surviving non-stem cancer cells can partially replenish the CSC compartment, thereby limiting the overall antitumor efficacy of radiation therapy. Dissecting and understanding the molecular mechanisms underlying this process has the potential to identify novel therapeutic strategies to target GSCs and improve patient outcomes.

Exosomes, have emerged as key mediators of intercellular communication within the tumor microenvironment, facilitating the transfer of oncogenic cargo, including proteins, nucleic acids, and lipids, between tumor cells and surrounding stromal cells [32].

In this study, we investigated the bi-directional crosstalk between GSCs and non-stem glioma cells mediated by exosomes. Treatment of differentiated GBM cells with gliomasphere-derived exosomes impaired self-renewal and significantly reduced functional GSC frequency. Previous studies have indicated that gliomasphere formation correlates with tumorigenicity and clinical outcome in GBM [33]. Consistent with this observation, scRNA-seq analysis demonstrated that gliomasphere-derived exosomes attenuated the radiation-induced increase of the NPC-like population, a cell population with known GSC properties [34, 35].

Functional annotation of the gliomasphere-derived exosomal cargo revealed strong enrichment of proteins involved in ribosomal function [36], mRNA translation[37–39], and RNA binding (RPL/RPS, EIF, EEF, PABPC1, SYNCRIP, HNRNPU) [39]. Given that these exosomes reduced both NPC-like cell abundance and stem-cell frequency following irradiation, their cargo profile is consistent with the modulation of translational and RNA regulatory pathways that restrain radiation-induced acquisition of the NPC-like state.

Furthermore, gliomasphere-derived exosomes were enriched for proteins involved in actin-remodeling, membrane–cytoskeletal, adhesion, and trafficking networks that normally support glioma cell motility. Despite the enrichment of this migration-associated machinery, gliomasphere derived exosomes significantly suppressed radiation-induced migration, indicating that the abundance of specific proteins in EV cargo does not necessarily reflect the activation of the corresponding pathways in recipient cells but may instead reflect the highly infiltrative nature of cells within gliomaspheres. The inhibitory phenotype resulting from exosome treatment of irradiated differentiated cells may instead arise from altered stoichiometry, localization, or regulation of cytoskeletal proteins following EV uptake, compromised chemokine sensing and chemotaxis, or from additional exosomal cargo that counteracts radiation-induced migratory signaling.

Only 17 DAPs were identified in exosomes of monolayer cells. The exosomal enrichment of NID2 and laminin subunits LAMA5, LAMB1, LAMB2 and LAMC1 indicates a basement-membrane-associated extracellular-matrix signature [40]. LAMA5 is of particular interest because laminin α5 is a structural basement-membrane component and has been shown to promote glioblastoma-cell proliferation and adhesion [41, 42], whereas the contribution of NID2, LAMB1, LAMB2 and LAMC1 to GSC expansion remains to be established directly.

IQGAP3 (IQ Motif Containing GTPase Activating Protein 3) and APOE are proliferation/metabolic signals. IQGAP3 is of particular interest as it was recently described as driver of GSC renewal and radioresistance through binding and stabilization of SOX2 [43].

The cell composition of exosome-treated remained largely untouched and the population of mixed-vascular-like cells increased only 1.13-fold. This cell population has been shown to contain rare neurovascular progenitor (NVP) cells that contribute to GBM heterogeneity and aggressiveness [20] but the moderate increase in NVP cells was insufficient to explain the observed increase in functional GSCs after exosome treatment. GSEA after pseudo-bulk analysis showed no positive enrichment of in stem cell-related gene sets in exosome-treated cells. Instead, exosome treatment was associated GSEA with positive enrichment in several gene sets associated with a more aggressive tumor phenotype, including HALLMARK_MYC_TARGETS_V1, HALLMARK_OXIDATIVE_PHOSPHORYLATION, and HALLMARK_E2F_TARGETS. These findings raise the possibility that the apparent increase in functional GSCs reflects the expression of pro-proliferative genes that enhance sphere-formation rather than true expansion of the stem cell population. Further studies are needed to determine whether exosome-treated gliomaspheres maintain elevated stem-cell numbers over time.

Together, these exosomal proteins suggest that differentiated glioma cells can promote GSC expansion by delivering a pro-stemness niche signal. The cargo is enriched for basement-membrane/ECM components and regulators (especially LAMA5, LAMB1/2, LAMC1, and NID2) that could enhance integrin-dependent adhesion, survival, and self-renewal signaling, while THBS1/TGFBI may further remodel the extracellular signaling environment and IQGAP3/APOE may support self-renewal, radioresistance, and/or metabolic states [43].

Overall, the data support a model in which exosomes from differentiated cells feed back onto the GSC compartment and increase gliomasphere stem-cell numbers by recreating or reinforcing a stem-cell-supportive ECM/signaling niche.

Our study has several limitations: First, we only focused on protein cargo of exosomes. It is possible that metabolite and/or RNA cargo contribute to the effect of exosomes on radiation-induced phenotype conversion or stem cell expansion. Especially, the high abundance of the RNA-binding protein SYNCRIP (Synaptotagmin-binding cytoplasmic RNA-interacting protein), also known as hnRNP Q (heterogeneous nuclear ribonucleoprotein Q) in exosomes derived from gliomaspheres indicate a possibility that the effects are mediated by miRNAs channeled specifically into exosomes by SYNCRIP [44, 45]. While sequencing the miRNA cargo was beyond the scope of this study, it warrants future investigation into miRNA-mediated suppressive effects on radiation-induced phenotype conversion. Lastly, our study was restricted to an *in vitro* setting and at this point it is unclear to which extend a normal human brain microenvironment will shape the cargo of tumor cell-derived exosomes.

### Conclusions

In this study, we identify exosomes as mediators of bidirectional communication between GSCs and non-stem glioma cells that regulates GBM cell-state homeostasis at steady state and following RT. GSC-derived exosomes suppressed RT-induced conversion of non-stem glioma cells into induced GICs, reduced functional GIC frequency, and prevented the radiation-associated expansion of NPC-like cells. Conversely, exosomes derived from non-stem glioma cells promoted GSC self-renewal and modestly increased the mixed-vascular-like population, which may contain neurovascular progenitor cells with stem-like properties. Proteomic analysis revealed distinct exosomal cargo profiles associated with these opposing effects, including enrichment of translational and cytoskeletal regulators in GSC-derived exosomes and extracellular-matrix and stem-cell-niche components in non-stem-cell-derived exosomes. Together, these findings support a model in which exosome-mediated feedback dynamically regulates the balance between stem-like and differentiated GBM cell states. Further studies are needed to identify the specific cargo responsible for these effects and determine whether disrupting this intercellular communication can improve therapeutic responses.

## Funding

FP was supported by grants from the *National Cancer Institute* (R01CA260886, R01CA281682). AB, HIK and FP were supported by the American Cancer Society (<u>CSCC-Team-23-980262-01-CSCC</u>).

## Author contributions

FP conceived of the study. LA and HH performed the experiments. HIK, LH, LML and AB provided materials. FP and LA analyzed the data and drafted the manuscript. All authors contributed to and approved the final version of the manuscript.

## Competing interests

The authors declare no conflict of interest

## Data and materials availability

All data are included in the article and/or *SI Appendix.* All cell lines will be made available upon reasonable request to the corresponding author. All sequencing data have been submitted to Gene Expression Omnibus and are available with the following Accession Number: GSE345825.

## Acknowledgements

This work was made possible, in part, through access to the following: The Electron Imaging Center for Nanosystems (EICN) at the University of California, Los Angeles’ California for NanoSystems Institute (CNSI) (RRID:SCR_022900) (NIH S10RR23057 and U24GM116792 to ZHZ). The Mass Spectrometry Facility at UCLA’s Molecular Instrumentation center. The UCLA Broad Stem Cell Research Center Flow Cytometry Core. The Genomics Research and Technology Hub (formerly Genomics High-Throughput Facility) Shared Resource of the Cancer Center Support Grant (P30CA-062203), the Single Cell Analysis Core shared resource of Complexity, Cooperation and Community in Cancer (U54CA217378), the Genomics-Bioinformatics Core of the Skin Biology Resource Based Center @ UCI (P30AR075047) at the University of California, Irvine and NIH shared instrumentation grants 1S10RR025496-01, 1S10OD010794-01, and 1S10OD021718-01.

---

**Supplementary Figure 1.**
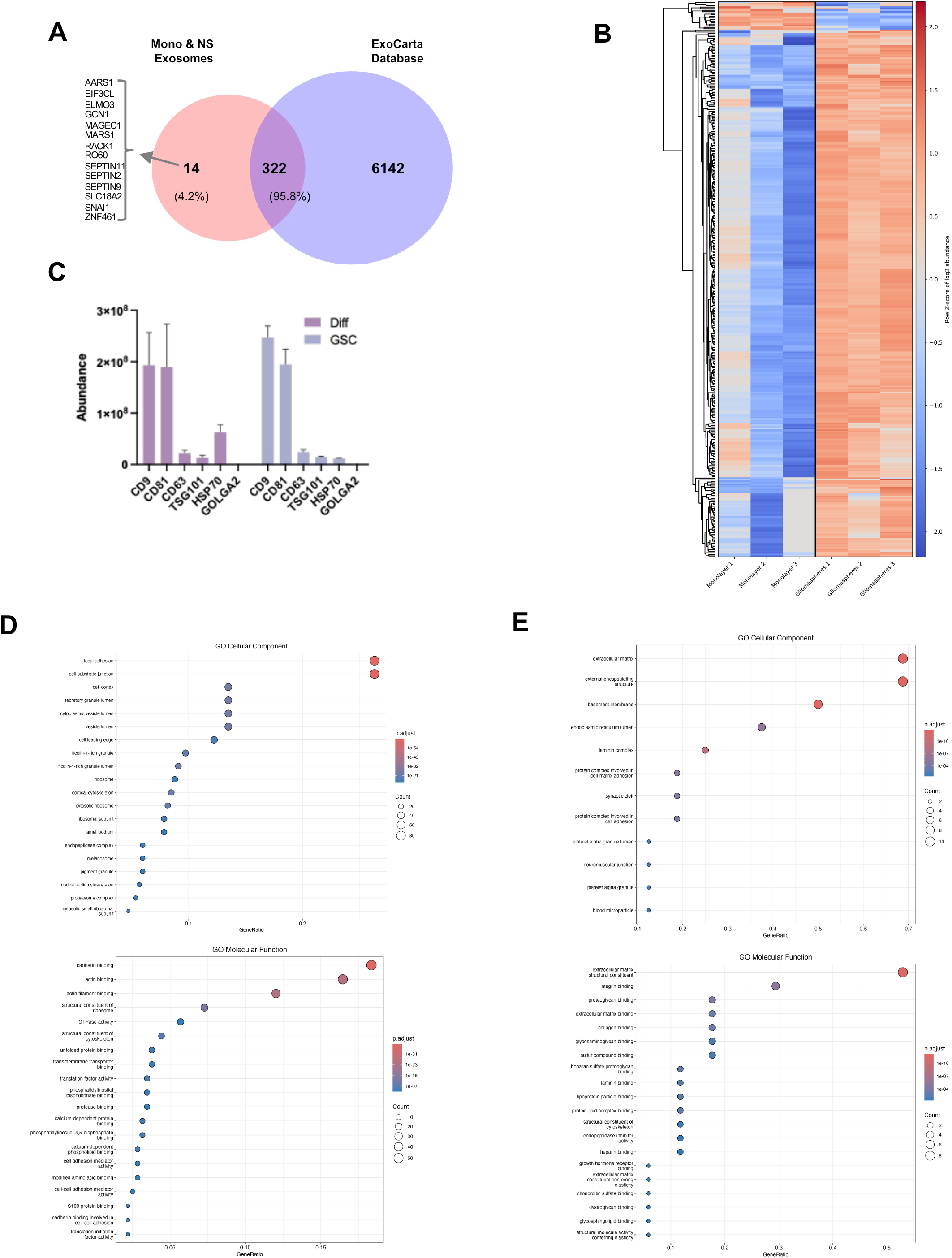
**A**) Venn diagram of top protein candidates in both gliomaspheres and monolayer EV samples against the ExoCarta database. **B**) Heatmap of z-scored protein intensities of differentially expressed top proteins (FDR <1%, p-value <0.05) across gliomasphere- and monolayer-derived EV samples (n=3 biological replicates per condition). **C**) Mass spectrometry-based abundance values for markers identified in exosomes from differentiated and gliomasphere samples. The Golgi marker GOLGA2 is used as a negative control to indicate successful enrichment of vesicles with no detectable Golgi protein contamination.

**Supplementary Figure 2.**
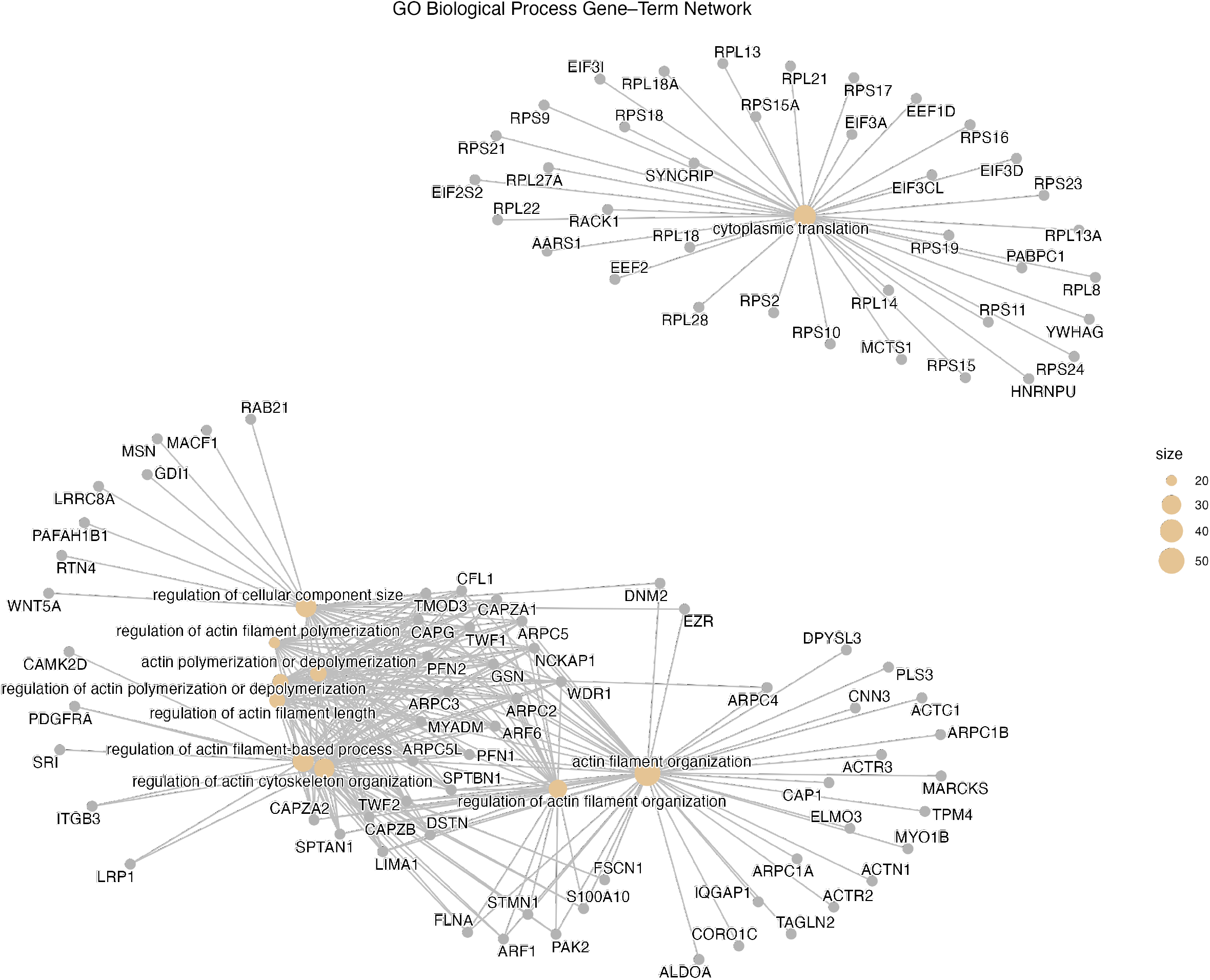
*Cytoplasmic Translation* and *Actin Filament Organization* gene-term network plots for DAPs in gliomasphere-derived exosomes.

**Supplementary Figure 3.**
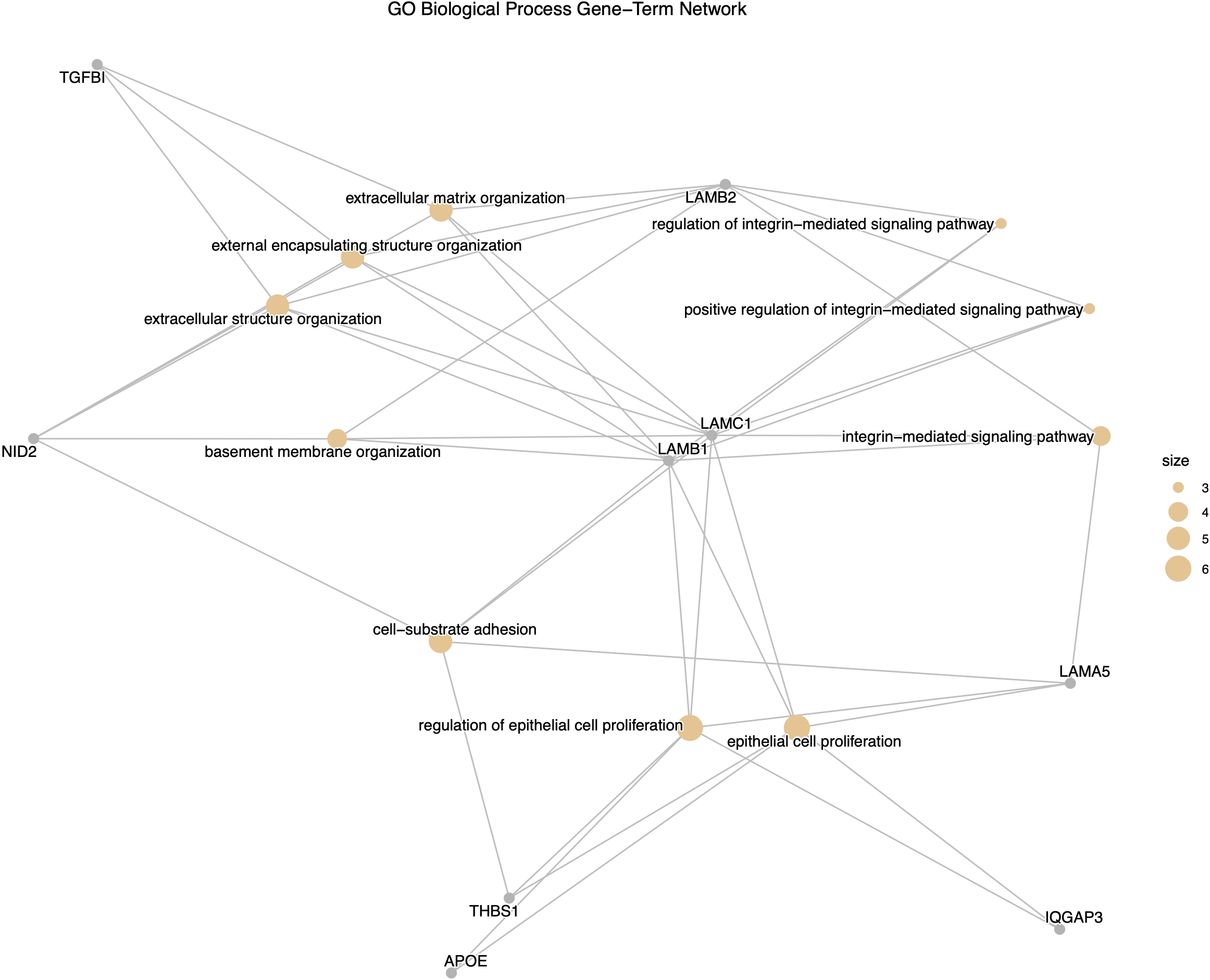
*G*ene-term network plots for DAPs in monolayer-derived exosomes

## Notes

### Competing Interest Statement

The authors have declared no competing interest.

